# Mitochondrial introgression by ancient admixture between two distant lacustrine fishes in Sulawesi Island

**DOI:** 10.1101/2020.12.29.424662

**Authors:** Mizuki Horoiwa, Ixchel F. Mandagi, Nobu Sutra, Javier Montenegro, Fadly Y. Tantu, Kawilarang W. A. Masengi, Atsushi J. Nagano, Junko Kusumi, Nina Yasuda, Kazunori Yamahira

## Abstract

Sulawesi, an island located in a biogeographical transition zone between Indomalaya and Australasia, is famous for its high levels of endemism. Ricefishes (family Adrianichthyidae) are an example of taxa that have uniquely diversified on this island. It was demonstrated that habitat fragmentation due to the Pliocene juxtaposition among tectonic subdivisions of this island was the primary factor that promoted their divergence; however, it is also equally probable that habitat fusions and resultant admixtures between phylogenetically distant species may have frequently occurred. Previous studies revealed that some individuals of *Oryzias sarasinorum* endemic to a tectonic lake in central Sulawesi have mitochondrial haplotypes that are similar to the haplotypes of *O. eversi*, which is a phylogenetically related but geologically distant (ca. 190 km apart) adrianichthyid endemic to a small lake. In this study, we tested if this reflects ancient admixture of *O. eversi* and *O. sarasinorum*. Population genomic analyses of genome-wide single-nucleotide polymorphisms revealed that *O. eversi* and *O. sarasinorum* are substantially reproductively isolated from each other. Comparison of demographic models revealed that the models assuming ancient admixture from *O. eversi* to *O. sarasinorum* was more supported than the models assuming no admixture; this supported the idea that the *O. eversi-like* mitochondrial haplotype in *O. sarasinorum* was introgressed from *O. eversi*. This study is the first to demonstrate ancient admixture of lacustrine organisms in Sulawesi beyond 100 km. The complex geological history of this island enabled such island-wide admixture of lacustrine organisms, which usually experience limited migration.

## Introduction

Sulawesi, an island located in a biogeographical transition zone between Indomalaya and Australasia, is famous for its high levels of endemism in both the terrestrial and freshwater fauna [1,2]. This endemism indicates that these taxa diversified within the island. Sulawesi is composed of three major tectonic subdivisions, two of which originated in the Asian and Australian continental margins, and the other emerged by the orogeny due to tectonic collision between the two plates [3–6]. These three tectonic subdivisions collided and have been juxtaposed with each other since the Pliocene (ca. 4 Mya), which resulted in the current shape of Sulawesi [7]. It was demonstrated that this complex geological history of the island may have largely affected the diversification of several Sulawesi endemic taxa (e.g., [8–11]).

Family Adrianichthyidae, commonly referred to as ricefishes or medaka, is one such taxon that has uniquely diversified on this island [11,12]. Previous studies revealed that adrianichthyids on this island are composed of six major clades (Fig 1) and demonstrated that divergence of these major clades largely reflected the tectonic activities of this island [11,13]. In particular, habitat fragmentation due to the Pliocene juxtaposition among the tectonic subdivisions was the primary factor that drove divergence of the lacustrine lineages distributed in tectonic lakes of central Sulawesi [11]. However, it is less known how species or populations within each clade have diverged.

**Fig 1.**
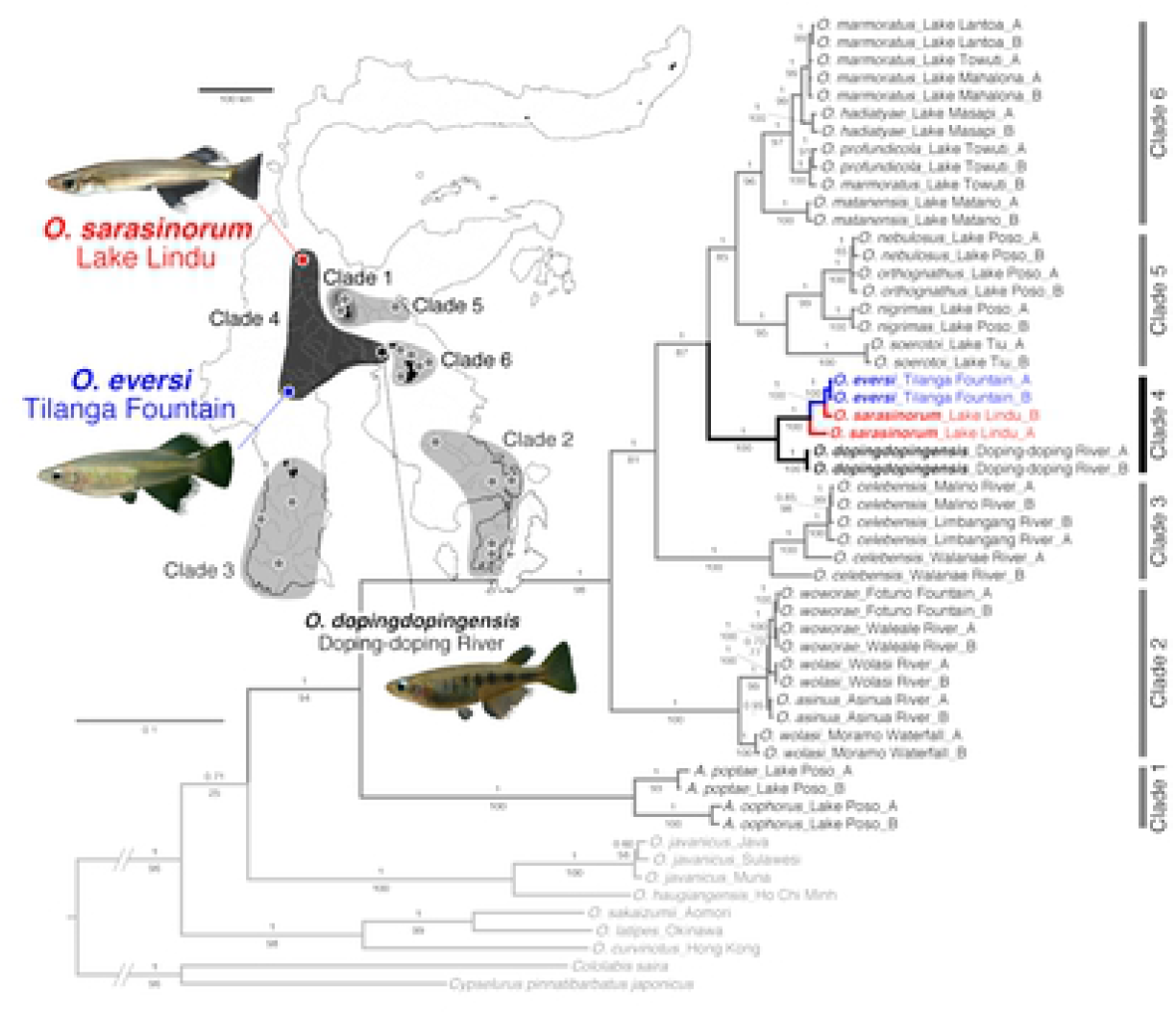
Mitochondrial phylogeny of Sulawesi adrianichthyids and a map of Sulawesi with the distribution of the major lineages. The mitochondrial phylogeny was based on cyt b (1,141 bp) and ND2 (1,046 bp) (modified from [13]). Note that *Oryzias sarasinorum* and *O. eversi* are endemic to Lake Lindu and Tilanga Fountain, respectively, which are approximately 190 km apart. Numbers on branches are Bayesian posterior probabilities (top) and maximum likelihood bootstrap values (bottom). The scale bar indicates the number of substitutions per site.

Within each major clade, individuals from a single species or population form a clade in most cases, which indicates that each species or population is phylogenetically distinct. However, there are several exceptions. For example, *O. sarasinorum, O. eversi*, and *O. dopingdopingensis*, which are endemic to Lake Lindu, Tilanga Fountain, and Doping-doping River, respectively, in western to central Sulawesi, form a major clade in mitochondrial phylogenies (named Clade 4 by [11,13,14]); however, two *O. sarasinorum* mitochondrial haplotypes are paraphyletic, and one of them forms a clade with *O. eversi* haplotypes (Fig 1). It remains unknown why these two mitochondrial haplotypes coexist in the *O. sarasinorum* population.

One possibility is mitochondrial introgression from *O. eversi* to *O. sarasinorum*. It is possible that the Pliocene juxtaposition of tectonic subdivisions of this island caused both fragmentations and fusions of tectonic lakes in central Sulawesi, which may have led to repeated isolations and admixtures of lacustrine organisms. Indeed, recent studies demonstrated ancient admixtures between lacustrine species that inhabit different lakes that are currently distant from each other in central Sulawesi [15]. It is possible that similar ancient admixture might have occurred between Lake Lindu and Tilanga Fountain that might have caused mitochondrial introgression.

Another possibility is incomplete lineage sorting (ILS). The topology of the mitochondrial tree might be incongruent with the topology of species tree because of ILS. ILS is very likely if *O. sarasinorum* and *O. eversi* are still young species that did not diverge a long time ago. To test if the paraphyly of *O. sarasinorum* mitochondrial haplotypes can be explained by ancient admixture versus ILS, comparisons of predefined models assuming different demographic histories by coalescent simulations (e.g., [16–18]) are very useful.

In this study, we first reconfirmed the composition of *O. sarasinorum* and *O. eversi* mitochondrial haplotypes by increasing the number of individuals examined. Second, we examined population genetic structures of the two species using genomewide single nuclear polymorphisms (SNPs). Third, we tested whether ancient admixture or ILS was more likely to explain the coexistence of two mitochondrial haplotypes within *O. sarasinorum* by coalescent-based demographic comparisons. Based on these results, we demonstrated that the two distinct mitochondrial haplotypes within *O. sarasinorum* reflect historical introgressive hybridization.

## Materials and Methods

### Field collections

Using a beach seine, we collected 10 juveniles each of *O. sarasinorum* and *O. eversi* from Lake Lindu (S01°20′02″, E120°03′09″) and Tilanga Fountain (S03°02′07″, E119°53′14″), respectively. They were preserved in 99% ethanol after being euthanized with MS-222. Total DNA was extracted from muscles of each of the 20 individuals using a DNeasy Blood & Tissue Kit (Qiagen, Hilden, Germany). Field collections were conducted with permission from the Ministry of Research, Technology, and Higher Education, Republic of Indonesia (research permit numbers 394/SIP/FRP/SM/XI/2014, 106/SIP/FRP/E5/Dit.KI/IV/2018, and 20/E5/E5.4/SIP.EXT/2019). We followed the Regulation for Animal Experiments at University of the Ryukyus for handling fishes, and all experiments were approved by the Animal Care Committee of University of the Ryukyus(2018099 and 2019084)

### Mitochondrial sequencing

The mitochondrial NADH dehydrogenase subunit 2 (ND2) gene was amplified for each of the 20 individuals (10 *O. sarasinorum* and 10 *O. eversi*) by PCR and Sanger sequenced using the methods and primers described by [11]. In addition, ND2 sequences of 10 *O. dopingdopingensis* individuals were retrieved from the DNA Data Bank of Japan (DDBJ) (LC551957–LC551966). *Oryzias dopingdopingensis* is a congener endemic to Doping-doping River in central Sulawesi (Fig 1). All sequences were aligned using the ClustalW option in MEGAX 10.1.8 [19], and the alignment was later manually corrected. We finally obtained 1,053 bp sequences of ND2 for the 30 individuals. The ND2 sequences of the 10 *O. sarasinorum* and 10 *O. eversi* individuals were deposited in DDBJ under accession numbers LC594688–LC594707.

### ddRAD sequencing

For the *O. sarasinorum* and *O. eversi* individuals, genomic data were generated by ddRAD-seq [20] (see [15] for details of library preparations). The library was sequenced with 50-bp single-end reads on an Illumina HiSeq 2500 system (Illumina, San Diego, USA) by Macrogen Japan Corporation (Kyoto, Japan). The sequencing reads were deposited in the DDBJ Sequence Read Archive under the accession number DRA011122. In addition, raw ddRAD-seq reads (50-bp single-end) for each of the 10 *O. dopingdopingensis* individuals were obtained from the DDBJ Sequence Read Archive (DRA010303).

Sequence trimming and mapping to a genome assembly of an *O. celebensis* individual (Ansai et al., unpublished data) were performed as described in [15]. Genotyping was conducted using the Stacks 1.48 software pipeline (*pstacks, cstacks*, and *sstacks*) [21,22] with default settings except for the minimum to create a “stack,” which was set to 10 reads (m = 10). The Stacks *populations* script was used to filter the loci that occurred in all three species (p = 3; i.e., *O. sarasinorum, O. eversi*, and *O. dopingdopingensis*) and in all individuals of each species (r = 1). Loci that deviated from Hardy–Weinberg equilibrium (5% significance level) in one or more species were excluded from the dataset using VCFtools 0.1.13 [23]. Genotype outputs were created in VCF format for only the first SNP per locus (write_single_SNP), which resulted in 1,487 SNP sites. In addition, a PHYLIP file of concatenated sequences was created (phylip_var_all), which resulted in 3,790 loci with a total length of 193,290 bp. Similarly, we created genotype outputs among only *O. sarasinorum* and *O. eversi* (i.e., p = 2) in VCF format, which resulted in 4,703 RAD loci that included 1,552 SNP sites.

### Phylogenetic analyses

A maximum-likelihood (ML) phylogeny among the 10 *O. sarasinorum*, 10 *O. eversi*, and 10 *O. dopingdopingensis* based on the 1,053-bp mitochondrial haplotypes was estimated with raxmlGUI 1.31 [24] using the codon-specific GTRGAMMAI model. *Oryzias dopingdopingensis* sequences were used as the outgroup, and bootstrap support values were calculated by a rapid bootstrap analysis of 1,000 bootstrap replicates. We also reconstructed a neighbor-joining (NJ) tree for the 193,290-bp concatenated RAD sequences using p-distances. Analysis was performed with MEGAX, and 1,000 bootstrap replicates were performed.

We also built a species tree based on the 1,487 RAD-seq SNPs using the Bayesian method implemented with SNAPP 1.4.1 [25]. Backward (U) and forward mutation rates (V) were estimated from the stationary allele frequencies in the data (U = 2.3066, V = 0.6384). Analysis was run using default priors with chainLength = 500,000 and storeEvery = 1,000. We discarded the first 10% of the trees as burn-in and visualized the posterior distribution of the remaining 450 trees as consensus trees using DensiTree 2.2.6 [26].

### Population structure analyses

We examined population structure within and among the three species with ADMIXTURE 1.3.0 [27] based on a PED file converted from the VCF file of the 1,487 RAD-seq SNPs using PLINK 1.90b4.6 [28]. ADMIXTURE was run for one to four clusters (i.e., K = 1–4). Statistical support for the different numbers of clusters was evaluated using the cross-validation technique implemented in ADMIXTURE. We also conducted principal component analyses using the R package SNPRelate 1.10.2 [29].

### Coalescence-based demographic inference

The demographic history of *O. sarasinorum* and *O. eversi* was inferred using fastsimcoal2 2.6.0.2 [17]. To better account for the complexity of multi-population models, we first compared five one-population models, which differ in population size change, separately for each species (S1 Fig) and chose the best-fit model for each species. One-hundred independent fastsimcoal2 runs with broad prior search ranges for each parameter were performed for each model using a one-dimensional site frequency spectrum created from the 4,703 RAD loci. We used a synonymous substitution rate of 3.5 × 10^-9^ per site and generation for each run, which was estimated using a cichlid parent–offspring trio with whole-genome sequencing [30] to convert the inferred parameters into demographic units. The relative fit of each model to the data was evaluated by Akaike information criterion (AIC) after transforming the log10-likelihood values to ln-likelihoods. As a result, the model incorporating population growth in the past (Pastgrowth_model) had the highest support in both species (S1 Table).

Next, we designed three types of two-population models using the best one-population model (Fig 2, S2 Table). The first type assumed allopatric divergence without gene flow and admixture (ALD_model). If this model was supported, the paraphyly of *O. sarasinorum* mitochondrial haplotypes would indicate ILS. The second and third types assumed gene flow (DGF_model) and ancient admixture (ADM_models), respectively. The third type of models was further divided into two, one of which assumed direct admixture between *O. eversi* and *O. sarasinorum* (ADM1_model) and the other assumed admixture between a lineage diverged from *O. eversi* and *O. sarasinorum* (ADM2_model). If DGF_model or ADM_models are supported, the scenario of mitochondrial introgression is highly probable. One-hundred independent fastsimcoal2 runs were performed for each model using a two-dimensional joint minor allele site frequency spectrum created from the 4,703 RAD loci and the substitution rate of 3.5 × 10^-9^ per site and generation. The relative fit of each model to the data was evaluated by AIC, as described above. For the best-fit model, 95% confidence intervals were calculated by parametric bootstrapping according to the program manual.

**Fig 2.**
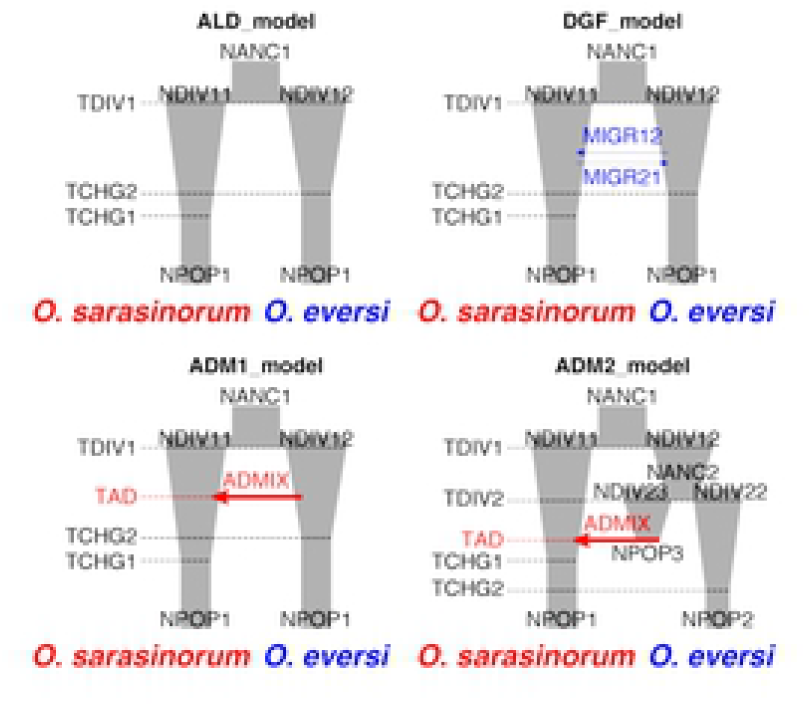
Schematic illustration of two-population demographic models. Note that growth was modeled as exponential and not linear as depicted here.

## Results

### Phylogeny and population structure

The mitochondrial ML phylogeny revealed two haplotype types within *O. sarasinorum* (Fig 3A), one of which was clustered with *O. eversi* haplotypes. The monophyly of this haplotype and *O. eversi* haplotypes had 100% ML bootstrap support. The other *O. sarasinorum* haplotypes formed a clade with 100% ML bootstrap support. In contrast, the nuclear NJ phylogeny based on the concatenated RAD sequences (193,290 bp) did not reveal two clusters within *O. sarasinorum* (Fig 3B). All *O. sarasinorum* individuals formed a clade with 99% bootstrap support. All *O. eversi* individuals also formed a clade with 99% bootstrap support. The species tree estimated by SNAPP also yielded the same topology (S2 Fig). In the posterior distribution of the species trees, all of the trees supported a topology consistent with the NJ tree.

**Fig 3.**
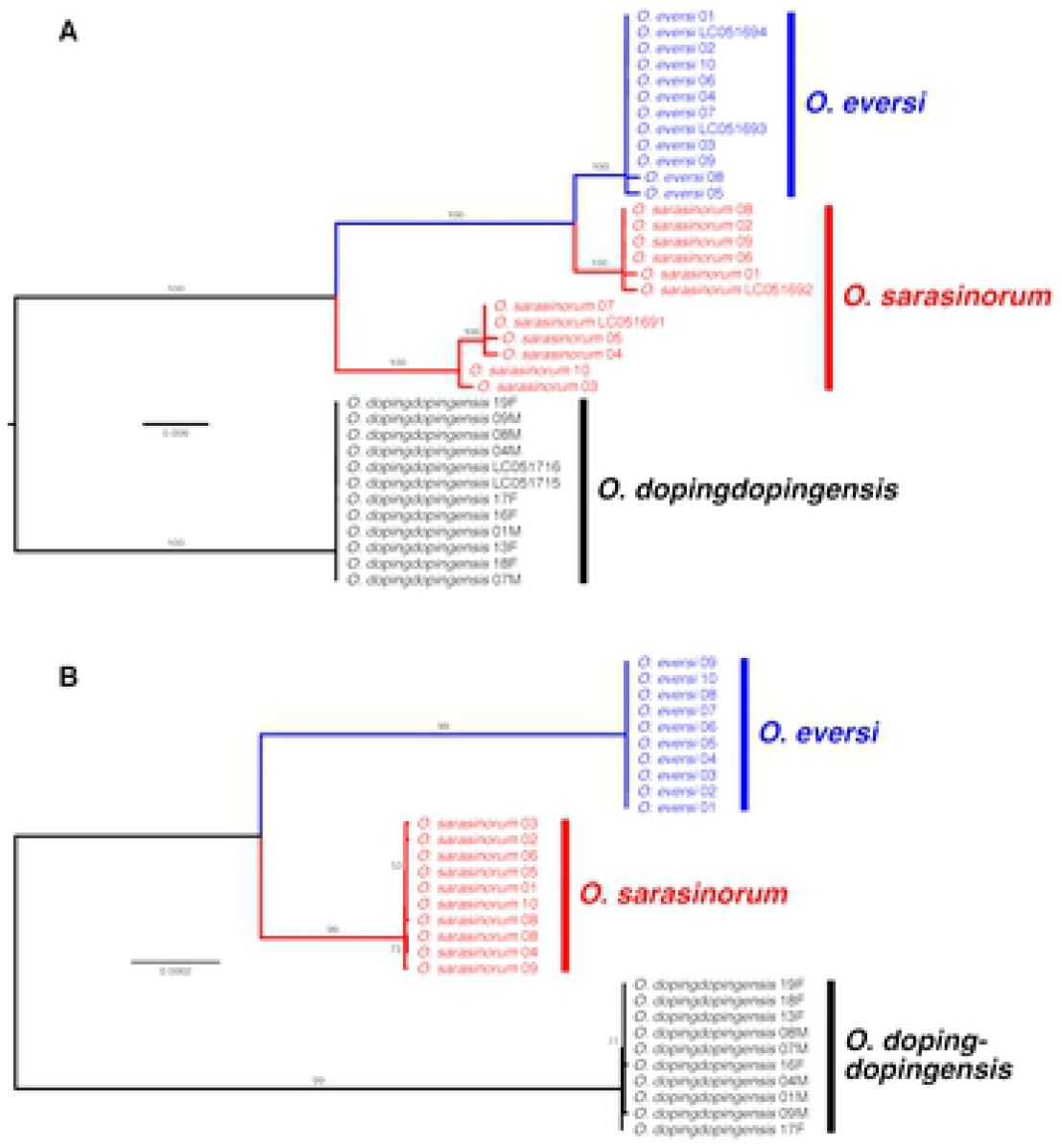
Phylogenies of *Oryzias sarasinorum* and *O. eversi*. (A) Maximum-likelihood phylogeny based on the 1,053-bp mitochondrial ND2 sequences and (B) neighborjoining phylogeny based on the 193,290-bp concatenated RAD sequences. Numbers on branches are bootstrap values.

ADMIXTURE analysis based on 1,487 SNPs revealed that the occurrence of three clusters (K = 3) had the highest support, and that *O. sarasinorum* and *O. eversi* were clearly separated (Fig 4). These two species were also separated from each other by the second principal component (PC2) in the principal component analysis (S3 Fig).

**Fig 4.**
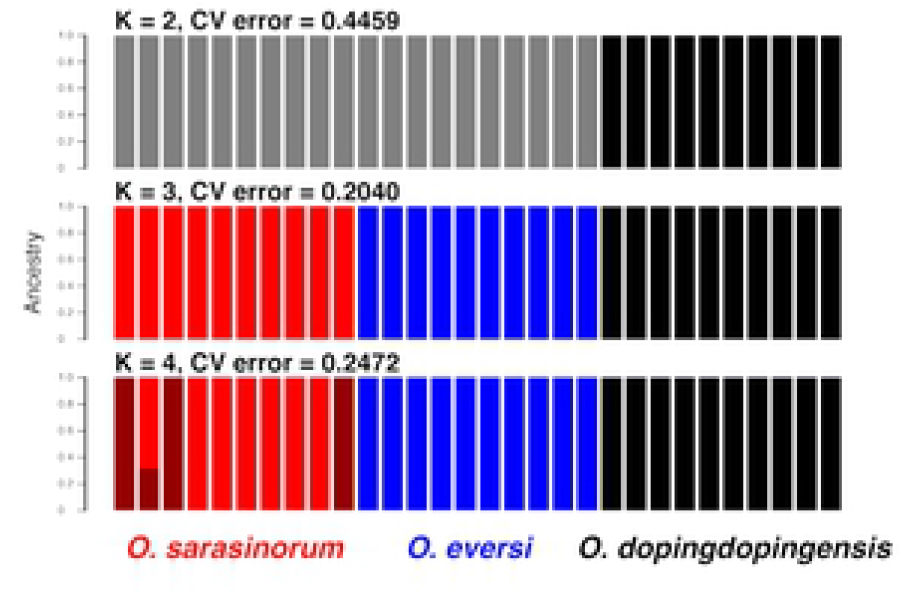
ADMIXTURE results showing K = 2–4 genetic clusters. Analysis was based on 1,487 SNPs among the three species.

### Demographic model selection

The model assuming direct ancient admixture (ADM1_model) was best supported by the fastsimcoal2 runs (Table 1). In this model, the common ancestor of *O. sarasinorum* and *O. eversi* diverged approximately 85,000 (78,867–158,420) generations ago (Fig 5A, Table 2). Population size of *O. sarasinorum* and *O. eversi* was estimated to have grown and shrunk, respectively, after they diverged from each other. Approximately 7,700 (1,898–21,372) generations ago, *O. sarasinorum* experienced introgression from *O. eversi*. The ratio of *O. eversi* migrants to *O. sarasinorum* was estimated to be 2.3% (1.2–6.1%).

**Fig 5.**
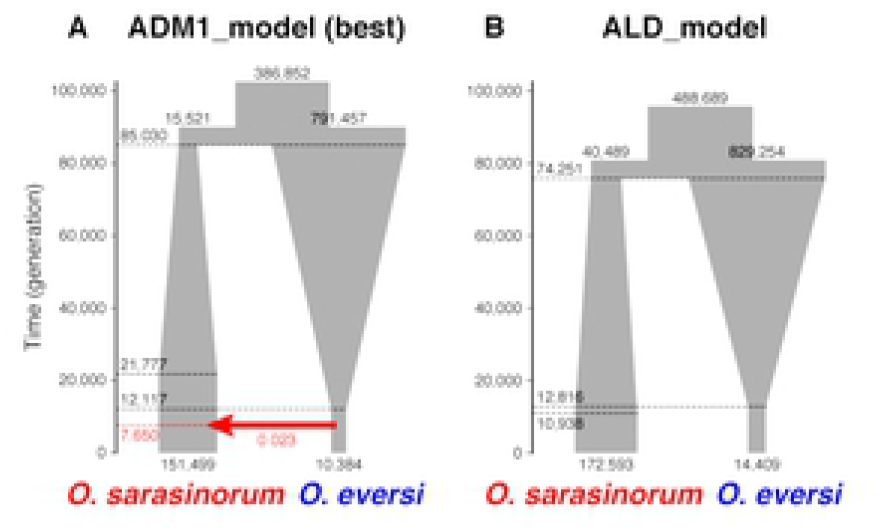
Schematic illustration of (A) ADM1_model (the best model) and (B) ALD_model estimated by fastsimcoal2 runs. The model is drawn to scale (time in generations) and population sizes; however, growth was modeled to be exponential and not linear as depicted here. The red arrow represents admixture.

**Table 1.** Support for each two-population model.

| Model | Number of parameters | log10-likelihood | Relative likelihood | ln-likelihood | AIC | Δ-AIC |
| --- | --- | --- | --- | --- | --- | --- |
| ADM1 | 10 | −6,390.633 | — | −14,714.976 | 29,449.953 | — |
| ADM2 | 15 | −6,390.714 | $8.299 \times 10^{-1}$ | −14,715.163 | 29,460.326 | 10.373 |
| DGF | 10 | −6,407.783 | $7.079 \times 10^{-18}$ | −14,754.466 | 29,528.931 | 78.979 |
| ALD | 8 | −6,410.675 | $9.078 \times 10^{-21}$ | −14,761.125 | 29,538.249 | 88.297 |

**Table 2.** Confidence intervals for each parameter in the best model (ADM1_model). The 95% confidence intervals were obtained from nonparametric bootstrapping.

| Parameter | 95% Confidence interval |
| --- | --- |
| NPOP1 | 111,545–196,162 |
| NPOP2 | 10,972–21,694 |
| NDIV11 | 11,714–605,060 |
| NDIV12 | 203,803–1,570,705 |
| NANC1 | 238,169–612,834 |
| TCHG1 | 4,542–154,938 |
| TCHG2 | 4,820–32,347 |
| TDIV1 | 78,867–158,420 |
| TAD | 1,898–21,372 |
| ADMIX | 0.01269–0.06063 |

The model assuming admixture between a lineage diverged from *O. eversi* and *O. sarasinorum* (ADM2_model) was second best (Table 1). The time of admixture (TAD = ca. 6,000 generations ago) and the ratio of the migrants (ADMIX = 4.3%) were estimated to be similar with those estimated by ADM1_model (S3 Table). These admixture models were much better supported than the model assuming gene flow (DGF_model) and the model assuming allopatric divergence with no gene flow and no admixture (ALD_model) (Table 1, Fig 5B, S3 Table).

## Discussion

### Ancient admixture and introgressive hybridization between the two distant lacustrine species

The mitochondrial phylogeny in this study revealed that *O. sarasinorum* mitochondrial haplotypes were not monophyletic, and some haplotypes were clustered with *O. eversi* haplotypes. However, the nuclear phylogeny showed monophyly of the *O. sarasinorum* individuals, which were clearly separated from *O. eversi* individuals. The population structure analyses also revealed that *O. sarasinorum* and *O. eversi* were clearly distinct from each other. These findings indicate that the two species are currently reproductively isolated from each other. However, the coalescence-based demographic analyses supported the scenario that assumes ancient admixture from *O. eversi* to *O. sarasinorum*; this indicates that the *O. sarasinorum* mitochondrial haplotypes that are close to those of *O. eversi* reflect introgression from *O. eversi* to *O. sarasinorum* rather than ILS.

Lake Lindu and Tilanga Fountain are currently ca. 190 km from each other. It is thought that the common ancestor of lacustrine adrianichthyids (Clades 4–6 in Fig 1) endemic to tectonic lakes in central Sulawesi was distributed in a big lake or lake system until the Pliocene (ca. 4 Mya), but it was later fragmented into several lakes or lake systems [11]. The sister relationship between *O. sarasinorum* and *O. eversi* indicates that there was a time when their common ancestor was isolated in a lake that was later divided into two smaller lakes: one is present-day Lake Lindu and the other Tilanga Fountain.

However, it is possible that the lake did not just undergo division. Some tectonic lakes and lake systems are known to have undergone repeated fragmentations and fusions, which caused repeated isolations and admixtures of lacustrine organisms [15]. It is probable that Lake Lindu and Tilanga Fountain were repeatedly connected to each other even after being divided. A long rift valley created by the action of the Palu–Koro fault system is located in the north–south direction between Lake Lindu and Tilanga Fountain [3,31]; if there was a time when this rift valley was a rift valley lake, then Lake Lindu and Tilanga Fountain would not have been as isolated from each other as they are now.

Indeed, a fossil of an adrianichthyid species, †*Lithopoecilus brouweride*, was reported from this rift valley (Gimpoe Basin) in the Miocene (ca. 23.0–5.3 Mya) geological stratum [32,33]. †*Lithopoecilus* is morphologically intermediate between *Oryzias* and *Adrianichthys*, a larger adrianichthyid genus [32], just like *O. sarasinorum* [34]. Although the exact generation time for these species remains unknown, assuming a generation time of 2 years as in [15], our coalescent-base demographic inference estimated that their divergence was approximately 170,000 (158,000–317,000) years ago. Therefore, we think that †*L. brouweride* is the common ancestor of *O. sarasinorum* and *O. eversi*. Either way, it is certain that there was an adrianichthyid in between Lake Lindu and Tilanga Fountain, which is currently land. It is therefore possible that the two species that are presently 190 km apart underwent historical admixture.

Assuming a generation time of two years, the age of the admixture between *O. eversi* and *O. sarasinorum* was estimated to be ca. 4,000–43,000 years ago. However, this estimate may be too young. The divergence between the *O. eversi* mitochondrial haplotypes and the *O. eversi-like O. sarasinorum* haplotypes (i.e., p-distance = 0.807%) would have occurred ca. 260,000–322,000 years ago assuming a substitution rate of 2.5%–3.1% per million years [35], which has been used for divergence-time estimation of Sulawesi adrianichthyids [2,11,15]. This discrepancy may indicate that the synonymous substitution rate used in the demographic inference (i.e., 3.5 × 10^-9^ per site and generation) was too high.

### Endemism shaped by island-wide admixture

In summary, we demonstrated that *O. sarasinorum* and *O. eversi* have a history of being admixed even though they are currently distributed in geologically distant tectonic lakes. Ancient admixture between adjacent lakes was recently demonstrated from other lakes and lake systems in central Sulawesi [15]; however, this study is the first to demonstrate admixture beyond 100 km. It is the geological history of Sulawesi that enabled such an island-wide admixture event of lacustrine organisms, which usually experience limited migration. We also think that such repeated admixtures may have promoted diversification of this freshwater fish group and probably other freshwater taxa, because it has been recognized that hybridization facilitates rapid speciation and adaptive radiation (e.g., [36–38]). The high levels of endemism in many terrestrial and freshwater fauna on Sulawesi may have been shaped by repeated admixture between distant lineages caused by the complex geological history of this island.

## Acknowledgments

We thank the Ministry of Research, Technology, and Higher Education, Republic of Indonesia (RISTEKDIKTI); the Faculty of Fisheries and Marine Science, Sam Ratulangi University; and the Faculty of Animal Husbandry and Fisheries, Tadulako University for the permit to conduct research in Sulawesi. We thank Mallory Eckstut, PhD, from Edanz Group (https://en-author-services.edanz.com/ac) for editing a draft of this manuscript.

## Supporting information

**S1 Fig. Schematic illustration of one-population demographic models.** Note that growth was modeled to be exponential and not linear as depicted here.

**S2 Fig. Species trees estimated by SNAPP based on 1,487 SNPs.** Thin lines represent individual species trees.

**S3 Fig. Principal component analysis of genetic variance based on 1,487 SNPs.**

**S1 Table. Support for one-population models defined in S1 Fig.**

**S2 Table. Explanation of each parameter used in the coalescent-based demographic inference.**

**S3 Table. Inferred maximum-likelihood parameters for each model.**

## Author Contributions

**Conceptualization:** Kazunori Yamahira

**Data curation:** Javier Montenegro, Kazunori Yamahira

**Formal analysis:** Mizuki Horoiwa, Nina Yasuda, Kazunori Yamahira

**Funding acquisition:** Junko Kusumi, Kazunori Yamahira

**Investigation:** Ixchel F. Mandagi, Nobu Sutra, Fadly Y. Tantu, Kawilarang W. A. Masengi, Atsushi J. Nagano, Kazunori Yamahira

**Methodology:** Junko Kusumi, Kazunori Yamahira

**Project administration:** Kazunori Yamahira

**Supervision:** Kazunori Yamahira

**Validation:** Kazunori Yamahira

**Visualization:** Kazunori Yamahira

**Writing – original draft:** Mizuki Horoiwa, Nina Yasuda, Kazunori Yamahira

**Writing – review & edition:** Junko Kusumi

## Data Availability Statement

The ND2 sequences obtained in this study were deposited in DDBJ under accession numbers LC594688–LC594707. The ddRAD-seq reads were deposited in the DDBJ Sequence Read Archive under accession number DRA011122.

## References

1 Whitten AJ, Mustafa M, Henderson GS. The ecology of Sulawesi. Singapore: Periplus; 2002.

2 Stelbrink B, Albrecht C, Hall R, von Rintelen T. The biogeography of Sulawesi revisited: is there evidence for a vicariant origin of taxa on Wallace’s “anomalous island”? Evolution 2012; 66:2252–2271. https://doi.org/10.1111/j.1558-5646.2012.01588.x PMID: 22759300

3 Moss SJ, Wilson EJ. Biogeographic implications of the tertiary palaeogeographic evolution of Sulawesi and Borneo. In: Hall R, Holloway JD, editors. Biogeography and geological evolution of SE Asia. Leiden: Backhuys Publishers; 1998. pp. 133–163.

4 Hall R. Continental growth at the Indonesian margins of southeast Asia. Arizona Geol Soc Digest 2009; 22: 245–258.

5 Hall R. Australia-SE Asia collision: plate tectonics and crustal flow. In: Hall R, Cottam MA, Wilson MEJ, editors. The SE Asian gateway: history and tectonics of Australia-Asia collision. London: The Geological Society of London; 2011. pp. 75–109.

6 Spakman W, Hall R. Surface deformation and slab-mantle interaction during Banda Arc subduction rollback. Nat Geosci 2010; 3: 562–566. https://doi.org/10.1038/ngeo917

7 Wilson ME, Moss SJ. Cenozoic paleogeographic evolution of Sulawesi and Borneo. Paleo3 1999; 145: 303–337. https://doi.org/10.1016/S0031-0182(98)00127-8

8 Evans BJ, Brown RM, McGuire JA, Supriatna J, Andayani N, Diesmos A, Iskandar DT, Melnick DJ, Cannatella DC. Phylogenetics of fanged frogs (Anura; Ranidae; *Limnonectesy*. testing biogeographical hypotheses at the Asian–Australian faunal zone interface. Syst Biol 2003; 52: 794–819. DOI:10.1080/10635150390251063 PMID: 14668118

9 Evans BJ, Supriatna J, Andayani N, Melnick DJ. Diversification of Sulawesi macaque monkeys: decoupled evolution of mitochondrial and autosomal DNA. Evolution 2003; 57: 1931–1946. https://doi.org/10.1111/j.0014-3820.2003.tb00599.x PMID: 14503633

10 von Rintelen T, Stelbrink B, Marwoto RM, Glaubrecht M. A snail perspective on the biogeography of Sulawesi, Indonesia: origin and intraisland dispersal of the viviparous freshwater gastropod *Tylomelania*. PLoS One 2014; 9: e98917. https://doi.org/10.1371/journal.pone.0098917 PMID: 24971564

11 Mokodongan DF, Yamahira K. Origin and intra-island diversification of Sulawesi endemic Adrianichthyidae. Mol Phylogenet Evol 2015; 93: 150–160. https://doi.org/10.1016/j.ympev.2015.07.024 PMID: 26256644

12 Takehana Y, Naruse K, Sakaizumi M. Molecular phylogeny of the medaka fishes genus *Oryzias* (Beloniformes: Adrianichthyidae) based on nuclear and mitochondrial DNA sequences. Mol Phylogenet Evol 2005; 36: 417–428. https://doi.org/10.1016/j.ympev.2005.01.016

13 Mokodongan DF, Yamahira K. Mitochondrial and nuclear phylogenies and divergence time estimations of Sulawesi endemic Adrianichthyidae. Data in Brief 2015; 5: 281–284. https://doi.org/10.1016/j.dib.2015.08.032 PMID: 26543892

14 Mandagi IF, Mokodongan DF, Tanaka R, Yamahira K. A new riverine ricefish of the genus *Oryzias* (Beloniformes, Adrianichthyidae) from Malili, central Sulawesi, Indonesia. Copeia 2018; 106: 297–304. https://doi.org/10.1643/CI-17-704

15 Sutra N, Kusumi J, Montenegro J, Kobayashi H, Fujimoto S, Masengi KWA, Nagano AJ, Toyoda A, Matsunami M, Kimura R, Yamahira K. Evidence for sympatric speciation in a Wallacean ancient lake. Evolution 2019; 73: 1898–1915. https://doi.org/10.1111/evo.13821 PMID: 31407798

16 Hey J, Nielsen R. Integration within the Felsenstein equation for improved Markov chain Monte Carlo methods in population genetics. Proc Natl Acad Sci USA 2007; 104: 2785–2790. https://doi.org/10.1073/pnas.0611164104 PMID: 17301231

17 Excoffier L, Dupanloup I, Huerta-Sánchez E, Sousa VC, Foll M. Robust demographic inference from genomic and SNP data. PLoS Genet 2013; 9: e1003905. https://doi.org/10.1371/journal.pgen.1003905 PMID: 24204310

18 Cornuet J-M, Pudlo P, Veyssier J, Dehne-Garcia A, Gautier M, Leblois R, Marin J-M, Estoup A. DIYABC v2.0: a software to make Approximate Bayesian Computation inferences about population history using Single Nucleotide Polymorphism, DNA sequence and microsatellite data. Bioinformatics 2014; 30: 1187–1189. https://doi.org/10.1093/bioinformatics/btt763 PMID: 24389659

19 Stecher G, Tamura K, Kumar S. Molecular Evolutionary Genetics Analysis (MEGA) for macOS. Mol Biol Evol 2020; 37: 1237–1239. https://doi.org/10.1093/molbev/msz312 PMID: 31904846

20 Peterson BBK, Weber JNJ, Kay EHE, Fisher HS, Hoekstra HE. Double digest radseq: an inexpensive method for de novo SNP discovery and genotyping in model and non-model species. PLoS One 2012; 7: e37135. https://doi.org/10.1371/journal.pone.0037135 PMID: 22675423

21 Catchen JM, Amores A, Hohenlohe PA, Cresko WA, Postlethwait J. Stacks: building and genotypng loci de novo from short-read sequences. G3–Genes Genom Genet 2011; 1: 171–182. https://doi.org/10.1534/g3.111.000240 PMID: 22384329

22 Catchen J, Hohenlohe PA, Bassham S, Amores A, Cresko WA. Stacks: an analysis tool set for population genomics. Mol Ecol 2013; 22: 3124–3140. https://doi.org/10.1111/mec.12354 PMID: 23701397

23 Danecek P, Auton A, Abecasis G, Albers CA, Banks E, DePristo MA, Handsaker RE, Lunter G, Marth GT, Sherry ST, McVean G, Durbin R, 1000 Genomes Project Analysis Group. The variant call format and VCFtools. Bioinformatics 2011; 27: 2156–2158. https://doi.org/10.1093/bioinformatics/btr330 PMID: 21653522

24 Silvestro D, Michalak I. RaxmlGUI: a graphical front-end for RAxML. Org Divers Evol 2012; 12: 335–337. https://doi.org/10.1007/s13127-011-0056-0

25 Bryant D, Bouckaert R, Felsenstein J, Rosenberg NA, RoyChoudhury A. Inferring species trees directly from biallelic genetic markers: bypassing gene trees in a full coalescent analysis. Mol Biol Evol 2012; 29: 1917–1932. https://doi.org/10.1093/molbev/mss086 PMID: 22422763

26 Bouckaert RR. DensiTree: making sense of sets of phylogenetic trees. Bioinformatics 2010; 26: 1372–1373.

27 Alexander DH, Novembre J, Lange K. Fast model-based estimation of ancestry in unrelated individuals. Genome Res 2009; 19: 1655–1664. http://www.genome.org/cgi/doi/10.1101/gr.094052.109. PMID: 19648217

28 Purcell S, Neale B, Todd-Brown K, Thomas L, Ferreira MAR, Bender D, Maller J, Sklar P, de Bakker PIW, Daly MJ, Sham PC. PLINK: a tool set for whole-genome association and population-based linkage analyses. Am J Hum Genet 2007; 81: 559–575. https://doi.org/10.1086/519795 PMID: 17701901

29 Zheng X, Levine D, Shen J, Gogarten SM, Laurie C, Weir BS. A high-performance computing toolset for relatedness and principal component analysis of SNP data. Bioinformatics 2012; 28: 3326–3328. https://doi.org/10.1093/bioinformatics/bts606 PMID: 23060615

30 Malinsky M, Svardal H, Tyers AM, Miska EA, Genner MJ, Turner GF, Durbin R. Whole-genome sequences of Malawi cichlids reveal multiple radiations interconnected by gene flow. Nat Ecol Evol 2018; 2: 1940–1955. https://doi.org/10.1038/s41559-018-0717-x PMID: 30455444

31 Nugrahaa AMS, Hall R. Late Cenozoic palaeogeography of Sulawesi, Indonesia. Palaeo3 2018; 490: 191–209. https://doi.org/10.1016/j.palaeo.2017.10.033

32 de Beaufort LF. On a fossil fish from Gimpoe (Central-Celebes). Verhandlungen Geologie en Mijnbouw Genoot. Nederland en Koloniën, Geology Series 1934; 10: 180–181.

33 Frickhinger KA. Fossilian atlas. Fische. Melle: Hans A. Bensch; 1991.

34 Parenti LR. A phylogenetic analyses and taxonomic revision of ricefishes, *Oryzias* and relatives (Beloniformes, Adrianichthyidae). Zool J Linn Soc 2008; 154: 494–610. https://doi.org/10.1111/j.1096-3642.2008.00417.x

35 Echelle AA, Carson EW, Echelle AF, Van Den Bussche RA, Dowling TE, Meyer A. Historical biogeography of the New-World pupfish genus *Cyprinodon* (Teleostei: Cyprinodontidae). Copeia 2005; 2005: 320–339. https://doi.org/10.1643/CG-03-093R3

36 Abbott R, Albach D, Ansell S, Arntzen JW, Baird SJE, et al. Hybridization and speciation. J Evol Biol 2013; 26: 229–246. https://doi.org/10.1111/j.1420-9101.2012.02599.x PMID: 23323997

37 Meier JI, Marques DA, Mwaiko S, Wagner CE, Excoffier L, Seehausen O. Ancient hybridization fuels rapid cichlid fish adaptive radiations. Nat Commun 2017; 8: 14363. https://doi.org/10.1038/ncomms14363 PMID: 28186104

38 Marques DA, Meier JI, Seehausen O. A combination view on speciation and adaptive radiation. Trends Ecol Evol 2019; 34: 531–544. https://doi.org/10.1016/j.tree.2019.02.008 PMID: 30885412

